# Ruminant-associated *Listeria monocytogenes* isolates belong preferentially to dairy-related hypervirulent clones: a longitudinal study in 19 farms

**DOI:** 10.1101/2021.07.29.454412

**Authors:** Carla Palacios-Gorba, Alexandra Moura, Jesús Gomis, Alexandre Leclercq, Ángel Gómez-Martín, Hélène Bracq-Dieye, María L. Mocé, Nathalie Tessaud-Rita, Estrella Jiménez-Trigos, Guillaume Vales, Ángel García-Muñoz, Pierre Thouvenot, Empar García-Roselló, Marc Lecuit, Juan J. Quereda

## Abstract

The increasing prevalence of *Listeria monocytogenes* infections is a public health issue. Although studies have shown that ruminants constitute reservoirs of this foodborne pathogen, little is known about its epidemiology and genetic diversity within ruminant farms. Here we conducted a large-scale genomic and epidemiologic longitudinal study of *Listeria* spp. in dairy ruminants and their environments, comprising 19 farms monitored for three consecutive seasons (*N*=3251 samples). *L. innocua* was the most prevalent *Listeria* spp, followed by *L. monocytogenes*. *L. monocytogenes* was detected in 52.6% of farms (prevalence in feces samples 3.8%, in farm environment samples 2.5%) and more frequently in cattle (4.1%) and sheep (4.5%) than in goat farms (0.2%). Lineage I accounted for 69% of *L. monocytogenes* isolates. Among animal samples, the most prevalent sublineages (SL) and clonal complexes (CC) were SL1/CC1, SL219/CC4, SL26/CC26 and SL87/CC87, whereas SL666/CC666 was prevalent in environmental samples. 61 different *L. monocytogenes* CTs (cgMLST sequence types) were found, 17 of them (27.9%) common to different animals and/or surfaces within the same farms. *L. monocytogenes* prevalence was not affected by farm hygiene but by season: the overall prevalence of *L. monocytogenes* in cattle farms was higher during winter, and in sheep farms was higher during winter and spring. Cows in their second lactation had a higher probability of *L. monocytogenes* fecal shedding than other lactating cows. This study highlights that dairy farms constitute a reservoir for hypervirulent *L. monocytogenes* and the importance of continuous animal surveillance to reduce the burden of human listeriosis.

**IMPORTANCE:** *Listeria monocytogenes* is a bacterial pathogen responsible for listeriosis, the foodborne disease with the highest hospitalization and case-fatality rate. Despite increasing evidence that dairy products and ruminant farms are important reservoirs of *L. monocytogenes*, little is known about the epidemiology and genetic diversity of *Listeria* spp. within dairy ruminant farms. We report the largest *Listeria* spp. longitudinal study in individual domestic animals, and the first using whole-genome sequencing for a deep isolate characterization. Here, we show that domestic ruminants can be asymptomatic carriers of pathogenic *Listeria*, that *L. monocytogenes* fecal shedding is often intermittent, and that hypervirulent *L. monocytogenes* clones are overrepresented in dairy farms. Moreover, we uncover the effect of seasons and lactation number on the prevalence of *L. monocytogenes* in ruminants. Our study highlights the need for *Listeria* spp. monitoring in farm animals to control the spread of hypervirulent *L. monocytogenes* and reduce the burden of human listeriosis.

## INTRODUCTION

The genus *Listeria* currently includes 26 recognized species of ubiquitous small rod-shaped gram-positive bacteria (1, 2). Only two of these species, *L. monocytogenes* (*L. monocytogenes*) and *L. ivanovii*, are considered pathogens (3). *L. monocytogenes* is an important foodborne pathogen that can cause human and animal listeriosis, a severe invasive infection with high hospitalization and fatality rates in humans (20–30%) (4). In immunocompromised individuals and the elderly, listeriosis manifests mostly as septicemia and central nervous system (CNS) infections. In pregnant women, listeriosis can lead to fetal or neonatal complications (4, 5).

Domestic ruminants can become infected by *L. monocytogenes* through ingestion of contaminated silage (3), which can result in rhombencephalitis, septicemia, and abortion. Animals may also be asymptomatic carriers and shed the bacterium in their feces (6–11). In dairy ruminants, *L. monocytogenes* can be transmitted to bulk tank milk (BTM) from fecal or environmental contamination of the udder surface (9, 12–15), as a consequence of poor hygiene in the milking parlor or as a consequence of intramammary infection (14, 15). The prevalence of *L. monocytogenes* in BTM of dairy cow farms can range between 1.2 and 16% (9, 16–18) and contaminated milk poses several risks for producers and consumers, namely: (*i*) the development of listeriosis after consumption of raw milk contaminated products; (*ii*) the development of biofilms in the milking equipment that contributes to persistent contamination of the BTM and (*iii*) cross-contamination of dairy processing plants, pasteurized dairy products or other food associated environments (19–23). Fecal shedding of *L. monocytogenes* also poses a risk for inter-animal transmission in dairy farms and contamination of agricultural environments and raw vegetables at the pre-harvest stages (24).

*L. monocytogenes* population is heterogeneous and can be classified into lineages (25), PCR genoserogroups (26), clonal complexes (CCs or clones) and sequence types (STs) as defined by multilocus sequence typing (MLST) (27), and sublineages (SLs) and cgMLST types (CTs), as defined by core genome MLST (cgMLST) (28). *L. monocytogenes* genetic heterogeneity also reflects different pathogenic potential among *L. monocytogenes* isolates, with some clones being more frequently isolated from human (e.g. CC1, CC2, CC4, and CC6) (29, 30) and ruminants (e.g. CC1) (31) clinical cases. Despite increasing evidence that dairy products and ruminant farms are important reservoirs of *L. monocytogenes* (7–11, 30, 32–34), little is still known about the genetic diversity, transmission dynamics and persistence of pathogenic *L. monocytogenes* in farm environments.

The objectives of the present study were: (*i*) to determine the prevalence of *Listeria* spp. in individual dairy ruminants and the farm environment in Spanish farms by a longitudinal study design; (*ii*) to characterize the genetic diversity and population structure of *L. monocytogenes* in dairy farms using whole genome sequencing; and (*iii*) to understand the transmission of *L. monocytogenes* at the farm level and the risk factors (season, production hygiene, lactation number and the days in milk (DIM) of current lactation) that influence it.

## RESULTS

### Prevalence of *Listeria* spp. in dairy farms

A total of 3,251 samples were collected from 19 Spanish dairy farms over three consecutive seasons (Figure 1, Table S1): 2,081 from animals (2,080 feces and 1 CNS infection case) and 1,170 from the surrounding farm environment (195 feed, 390 food and water troughs, 195 beddings, 195 milk filters, and 195 milking station floor). Each farm was sampled one time per season except farm “Sheep B” which was subjected to 8 additional samplings from 2019 to 2020 (extra samples *n*=400; farm environment *n*=144; animal feces *n*=256, see M&M and Figure 1). None of the farms reported listeriosis cases, except one farm (“Sheep C”), where a listeriosis outbreak occurred on the last season sampled (Spring 2020).

**Figure 1.**
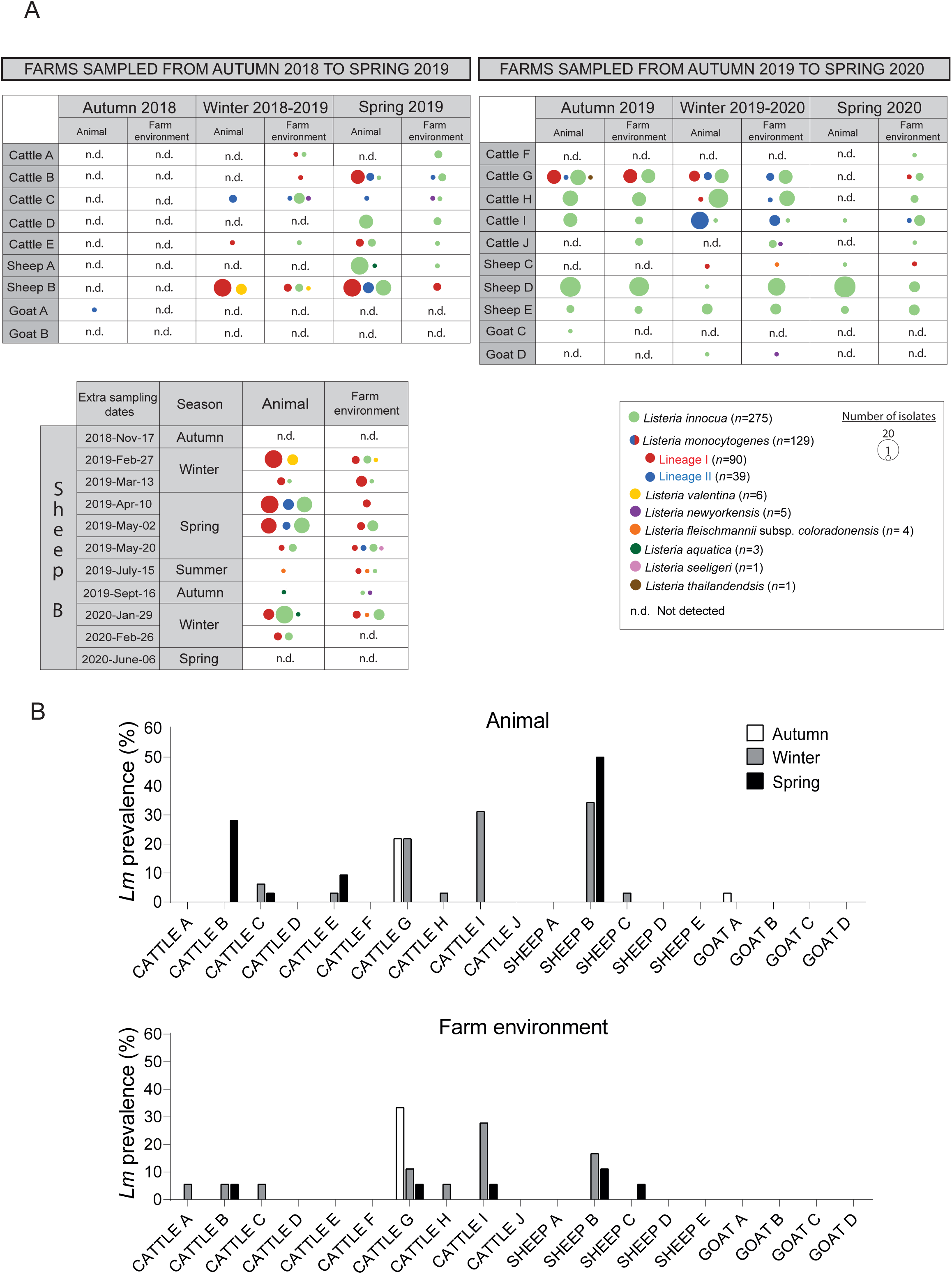
A) *Listeria* spp. isolated in this study from dairy cattle, sheep and goat farms during three consecutive seasons. Farm “Sheep B” was sampled 11 times during 7 consecutive seasons from autumn 2018 to spring 2020 (see Material and methods). Circle size is proportional to the number of isolates. B) Prevalence of *L. monocytogenes* in feces samples and the farm environment during three consecutive seasons. For consistency among farms, only data from three consecutive seasons (autumn 2018-Nov-07, winter 2019-Feb-27, and spring 2019-Apr-10) were considered for prevalence calculation on farm “Sheep B”.

*Listeria* spp. was detected in 94.7% (18/19) of farms and in all sampling seasons (Fig. 1). Overall, *Listeria* spp. prevalence was 11.2% (318/2850), and similar in feces samples (10.2%; 186/1824) and farm environment samples (12.9%; 132/1026) (Figure 1, Table S2). Prevalence varied significantly between farms from 0% to 43.3% and was overall higher in cattle and sheep farms (Table S2). The most prevalent species were *L. innocua* (64.7%; 275/425) and *L. monocytogenes* (30.6%; 130/425) (Table 1). Co-occurrence of the two *Listeria* species was detected in 0.8% (14/1824) of individual animal feces and 1.1% (11/1026) environmental samples (Table S3). *L. monocytogenes* was detected in 52.6% of farms (10/19), and prevalence in positive farms ranged between 0.7% and 21.3%, and was frequently higher for cattle farms (Fig. 1B, Table S2). *L. monocytogenes* prevalence was also comparable in feces samples (3.8%; 70/1824) and farm environment samples (2.5%; 26/1026), although in some farms, *L. monocytogenes* was most frequently present in feces samples compared to environmental samples in a specific season (Fig. 1B). *L. innocua* was present in 6.7% (123/1824) of feces samples and in 10.8% (111/1026) of farm environmental samples (Table S3). Among environmental samples, *L. monocytogenes* was detected in all sampled sites except in the milking station floor where other *Listeria* spp. were present (8 *L. innocua* and 1 *L. newyorkensis*). *L. monocytogenes* occurred on food troughs (5.8%), beddings (3.5%), feed (2.9%), milk filter socks (MFS) (2.3%), and the water troughs (0.6%) (Tables S2 and S3). Of note, no *L. monocytogenes* was detected in feed, food troughs and water troughs from 4 out of 9 of the farms where fecal shedders were detected.

**Table 1.** Number of isolates obtained in this study (*N*=425).

| *Listeria* species | Animal | Farm env. | Feed | Total |
| --- | --- | --- | --- | --- |
| *L. monocytogenes* | 89 (20.9%) | 32 (7.5%) | 9 (2.1%) | 130 (30.6%) |
| *L. innocua* | 150 (35.3%) | 92 (21.6%) | 33 (7.8%) | 275 (64.7%) |
| *L. valentina* | 5 (1.2%) | 1 (0.2%) |  | 6 (1.4%) |
| *L. newyorkensis* |  | 5 (1.2%) |  | 5 (1.2%) |
| *L. fleischmannii* subsp.<br>*coloradonensis* | 1 (0.2%) | 2 (0.5%) | 1 (0.2%) | 4 (0.9%) |
| *L. aquatica* | 3 (0.7%) |  |  | 3 (0.7%) |
| *L. seeligeri* |  | 1 (0.2%) |  | 1 (0.2%) |
| *L. thailandensis* | 1 (0.2%) |  |  | 1 (0.2%) |

Among animals, different seasonal patterns of *L. monocytogenes* shedding were observed (Fig. 1B). While no *L. monocytogenes* was detected in animals from farm “Cattle B” during autumn and winter, it increased sharply in the spring season (28.1%). Farm “Cattle G” was characterized by a high *L. monocytogenes* prevalence of fecal shedders during autumn and winter (21.9% in both seasons), but none was detected in spring. In farm “Cattle I”, although no *L. monocytogenes* was detected in autumn, it increased sharply to 31.2% in winter and disappeared in the spring sampling. Finally, while none of the tested sheep shed *L. monocytogenes* during autumn in farm “Sheep B”, the prevalence increased to 34.3% during winter and to 50% during spring (Fig. 1B).

Interestingly, in farm “Sheep C”, where an 8-week listeriosis outbreak occurred in the spring of 2020, no *L. monocytogenes* was isolated from any of the feces or environmental samples collected. The outbreak caused 89 abortions (from a total of 974 pregnant sheep) and CNS symptoms (inappetence, recumbency, difficulties in swallowing, drooping eyelid, ear and lip, head-tilt and circling) in 6 animals (1.6% total mortality). On samples taken *post-mortem* from the bedding surfaces, feces and the brainstem of one diseased sheep, *L. monocytogenes* was isolated from the brainstem and from one bedding sample but only *L. innocua* was detected on the feces (Table S3).

### Population structure and genetic diversity of *Listeria* spp. in dairy farms

Altogether, 425 *Listeria* spp. isolates were obtained from the farm environment (176/425, 41.4%) and animal samples (249/425, 58.6%) and further characterized at the genomic level. Eight different *Listeria* species were identified: *L. monocytogenes* (*n=*130, 30.6%)*, L. innocua* (*n=*275, 64.7%)*, L. valentina* (*n=*6, 1.4%), *L. newyorkensis* (*n=*5, 1.2%), *L. fleischmannii* subsp. *coloradonensis* (*n=*4, 0.9%)*, L. aquatica* (*n=*3; 0.7%)*, L. seeligeri* (*n=*1, 0.2%) and *L. thailandensis* (*n=*1, 0.2%) (Fig. 1, Fig. 2, and Table 1).

**Figure 2.**
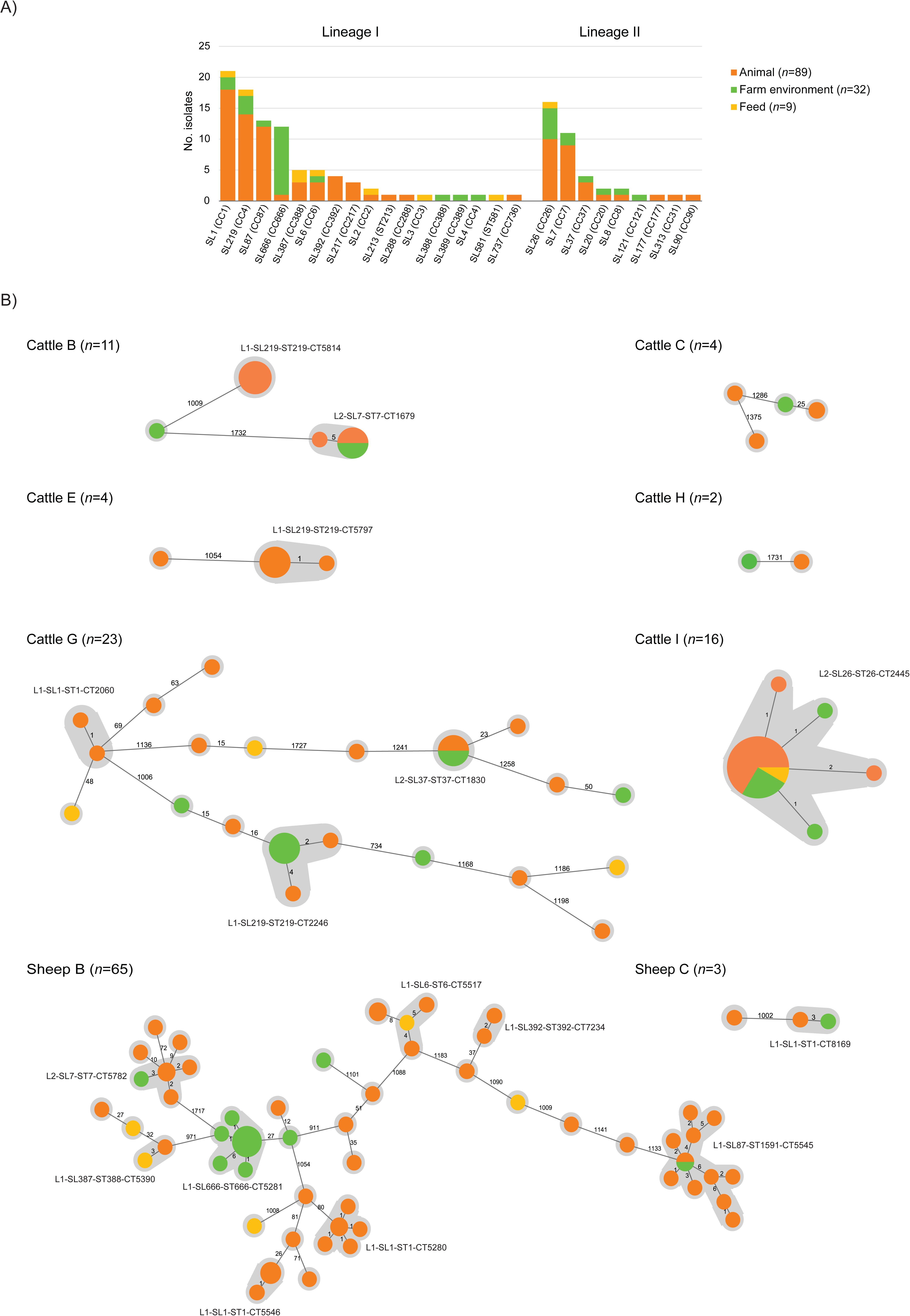
Diversity of *L. monocytogenes* isolates collected in this study (*n*=130) based on cgMLST (1748 loci) analyses. A) Distribution of sublineages in animals, feed and farm environmental sources. Corresponding clonal complexes (CCs), defined on the basis of 7-locus MLST, are shown in brackets. B) Minimum spanning tree based on the cgMLST profiles *L. monocytogenes* observed in each farm sampled in this study. Farms with only one *L. monocytogenes* isolate (“Cattle A” and “Goat A”) are omitted. Circle sizes are proportional to the number of isolates and are colored by source type, as in panel A. Values shown in connecting lines denote the number of allelic differences between profiles. Grey zones delimitate isolates within the same cgMLST type (cut-off of 7-allelic differences (28)) and types common to more than 1 isolate are labeled.

*L. monocytogenes* belonged to lineages I (*n=*91; 70%; genoserogroups IVb, *n*=70 and IIb, *n =* 21) and II (*n=*39; 30%; genoserogroup IIa) (Fig. 2 and Fig. 3). Among animal samples, the most prevalent SLs and CCs were SL1/CC1 (*n*=18, 13.8%), SL219/CC4 (*n=*14, 10.8%), SL26/CC26 (*n*=10, 7.7%) and SL87/CC87 (*n*=12, 9.2%), whereas SL666/CC666 (*n*=12, 9.2%) was prevalent in environmental samples (Fig. 2A, Table 2).

**Figure 3.**
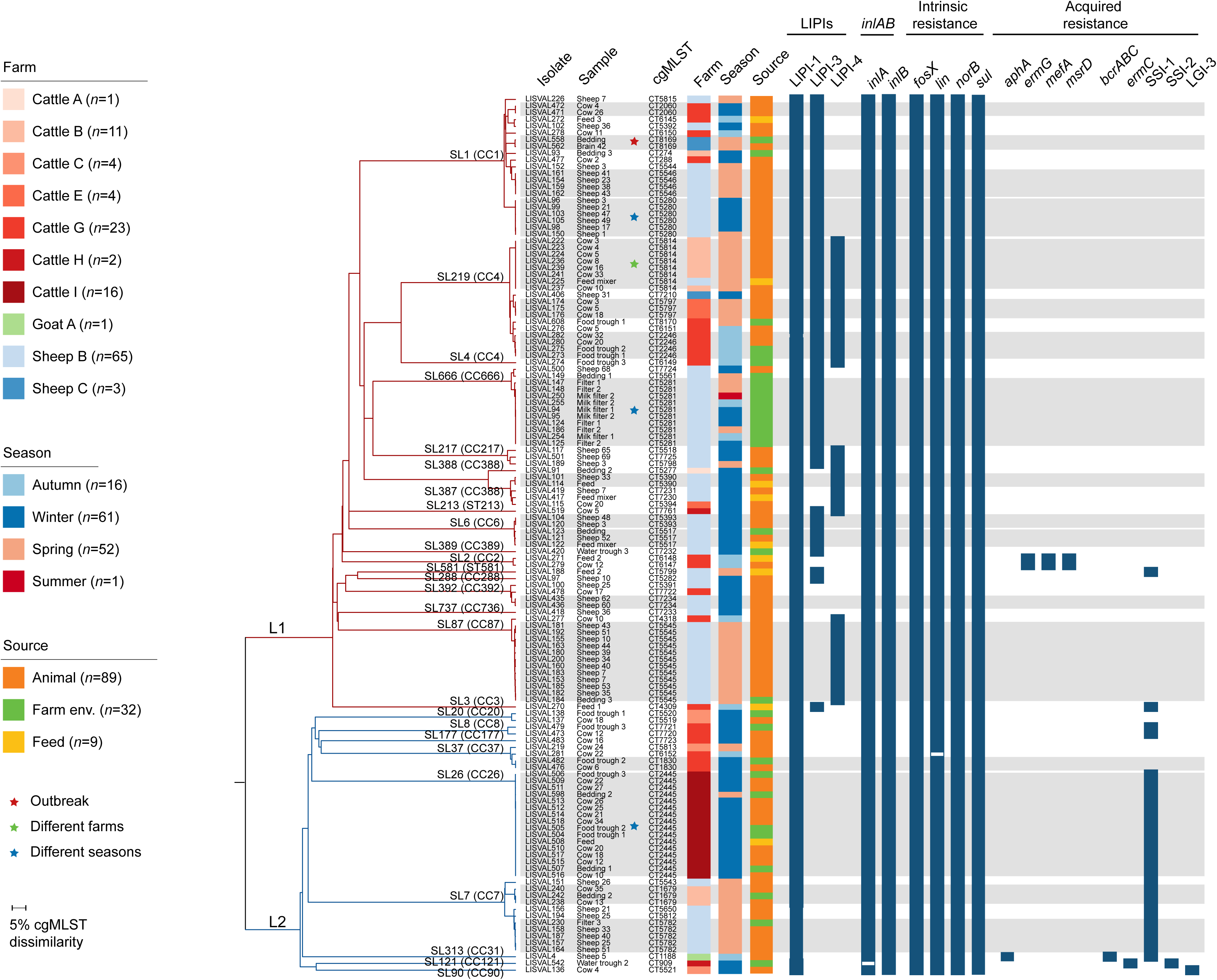
Genetic diversity of the 130 *L. monocytogenes* isolates sequenced in this study. A) cgMLST single linkage dendrogram. Branches are colored by phylogenetic lineage (L1, red; L2, blue) and labelled by sublineages. Corresponding clonal complexes (CCs), defined on the basis of 7-locus MLST, are shown in brackets. Isolates’ names, type of sample and cgMLST types are shown in tips. Colored boxes indicate the sampling farm, season and source, colored according with the key panel, respectively. Color-filled dark blue boxes indicate the presence of selected genetic traits: *Listeria* pathogenic islands (LIPI-1, LIPI-3 and LIPI-4), internalins (*inlA*, *inlB*), intrinsic antibiotic resistance (*fosX, lin, norB, sul*) and acquired resistance loci towards antibiotics (*aphA*, *ermG*, *mefA*, *msrD*), benzalkonium chloride (*bcrABC, ermC*), pH or oxidative stress (SSI-1, SSI-2) and metals (LGI-3). White-filled blue boxes represent genes with truncations leading to premature stop codons.

**Table 2.** *L. monocytogenes* cgMLST types detected comprising 2 or more isolates (*n*=17 types out of 61).

| cgMLST type | No. isolates | Farm(s) | Season(s) | Source(s) | Sources description | Traits |
| --- | --- | --- | --- | --- | --- | --- |
| *Lineage I* |  |  |  |  |  |  |
| L1-SL87-ST1591-CT5545 (CC87, IIb) | 12 | Sheep B | Spring 2019 (Apr, May) | A,E | feces (11), bedding (1) | LIPI-1,LIPI-4 |
| L1-SL666-ST666-CT5281 (CC666, IVb) | 10 | Sheep B | Winter 2019 (Feb, Mar), Spring 2019 (Apr, May), Summer 2019 (Jun), Autumn 2019 (Sep) | E | milk filter (10) | LIPI-1,LIPI-3 |
| L1-SL219-ST219-CT5814 (CC4, IVb) | 8 | Sheep B, Cattle B | Spring 2019 (May) | A,E | feces (7), feed (1) | LIPI-1,LIPI-3,LIPI-4 |
| L1-SL1-ST1-CT5280 (CC1, IVb) | 6 | Sheep B | Winter 2019 (Feb), Spring 2019 (Apr) | A | feces (6) | LIPI-1,LIPI-3 |
| L1-SL1-ST1-CT5546 (CC1, IVb) | 4 | Sheep B | Spring 2019 (Apr) | A | feces (4) | LIPI-1,LIPI-3 |
| L1-SL219-ST219-CT2246 (CC4, IVb) | 4 | Cattle G | Autumn 2019 (Oct) | A,E | feces (2), food trough (2) | LIPI-1,LIPI-3,LIPI-4 |
| L1-SL219-ST219-CT5797 (CC4, IVb) | 3 | Cattle E | Spring 2019 (Apr) | A | feces (3) | LIPI-1,LIPI-3,LIPI-4 |
| L1-SL6-ST6-CT5517 (CC6, IVb) | 3 | Sheep B | Winter 2019 (Mar) | A,E | feces (1), feed (1), bedding (1) | LIPI-1,LIPI-3 |
| L1-SL1-ST1-CT2060 (CC1, IVb) | 2 | Cattle G | Winter 2020 (Feb) | A | feces (2) | LIPI-1,LIPI-3 |
| L1-SL6-ST6-CT5393 (CC6, IVb) | 2 | Sheep B | Winter 2019 (Feb, Mar) | A | feces (2) | LIPI-1,LIPI-3 |
| L1-SL392-ST392-CT7234 (CC392, IIb) | 2 | Sheep B | Winter 2020 (Jan) | A | feces (2) | LIPI-1 |
| L1-SL387-ST388-CT5390 (CC388, IVb) | 2 | Sheep B | Winter 2019 (Feb) | A,E | feces (1), feed (1) | LIPI-1,LIPI-4 |
| L1-SL1-ST1-CT8169 (CC1, IVb) | 2 | Sheep C | Spring 2020 (May) | A,E | brain (1), bedding (1) | LIPI-1,LIPI-3 |
| *Lineage II* |  |  |  |  |  |  |
| L2-SL26-ST26-CT2445 (CC26, IIa) | 16 | Cattle I | Winter 2020 (Feb), Spring 2020 (May) | A,E | feces (10), feed (1), food trough (3), bedding (2) | LIPI-1, SSI-1 |
| L2-SL7-ST7-CT5782 (CC7, IIa) | 5 | Sheep B | Spring 2019 (Apr, May) | A,E | feces (4), milk filter (1) | LIPI-1, SSI-1 |
| L2-SL7-ST7-CT1679 (CC7, IIa) | 3 | Cattle B | Spring 2019 (May) | A,E | feces (2), bedding (1) | LIPI-1, SSI-1 |
| L2-SL37-ST37-CT1830 (CC37, IIa) | 2 | Cattle G | Winter 2020 (Feb) | A,E | feces (1), food trough (1) | LIPI-1 |

Sixty-one different cgMLST types were detected: 44 (72.1%) comprising only one isolate and 17 (27.9%) comprising 2 to 16 isolates sharing 0-6 allelic differences (Table 2, Fig. 2B). Up to 5 and 8 different CTs could be isolated from environmental and fecal materials, respectively, on a single sampling day (Table S3). Overall, a higher diversity was found in animal fecal samples than environmental samples (Shannon indexes 3.4 vs 2.8). The Hutcheson t-test showed a highly significant difference in terms of community composition of *L. monocytogenes* CTs (*P*=0.006), in the two categories (animal feces vs. environmental samples). No CTs were common to multiple farms, except for L1-SL219-ST219-CT5814 which was detected in farms “Sheep B” and “Cattle B”, separated by 73 km and sampled 7 days apart (Table S3). 16.4% (10/61) CTs were common to both environmental and animal samples from the same farm, 3 of them collected at different time points (Tables 2 and S3). Persistent *L. monocytogenes* strains (*i.e*. continued presence of a given CT over time, at a specific location (35)) were identified in 6 dairy farms (Table 2) and in one sheep (Tables S3 and S4).

*Listeria* pathogenic islands LIPI-3 and LIPI-4, where present in 50% (65/130) and 32.3% (42/130) *L. monocytogenes*, whereas all isolates carried LIPI-1, which is part of the *L. monocytogenes* core genome (Fig. 3). Acquired resistance traits towards antibiotics, disinfectants or other stress conditions, were rare (Table 3, Table S3 and Fig. 3). Two isolates (1.5%, cgMLST types L1-SL2-ST2-CT6147 and L1-SL2-ST2-CT6148) harbored genes conferring resistance to macrolides (*ermG*, *mefA* and *msrD* genes) or to benzalkonium chloride (*bcrABC* and Tn*6188::ermC;* cgMLST type L2-SL313-ST325-CT1188 and L2-SL121-ST121-CT909, respectively). SSI-1 (32/130 isolates, 24.6%, tolerance to low pH and high salt), SSI-2 (1/130, 0.7%; tolerance to alkaline and oxidative stress conditions) and LGI-3 (1/130, 0.7%; tolerance to cadmium) genomic regions were also present (Table 3). Acquired resistance genes were also present in *L. innocua* (19/275, 6.9%) and *L. aquatica* (1/3, 33.3%), with *tetM* (resistance to tetracyclines) being the most prevalent resistance trait detected (Table 3 and Table S3). The most prevalent CTs *L. monocytogenes* detected in this study (L1-SL87-ST1591-CT5545, *n*=12; L1-SL666-ST666-CT5281, *n*=10; L1-SL219-ST219-CT5814, *n*=8; and L2-SL26-ST26-CT2445, *n*=16) harbored at least one of the following genomic regions: LIPI-3, LIPI-4 or SSI-1 (Table 2, Fig. 3).

**Table 3.** Acquired antibiotic resistance genes detected in this study.

| *Listeria* species | No. isolates | Genes<sup>1</sup> | Farm(s) | Season(s) | Source(s) | Source description |
| --- | --- | --- | --- | --- | --- | --- |
| *L. monocytogenes* | 2 | *mefA,msrD,ermG* | Cattle G | Autumn 2019 (Oct) | A,E | feces (1), feed (1) |
|  | 1 | *aphA* | Goat A | Autumn 2018 (Nov) | A | feces (1) |
| *L. innocua* | 13 | *tetM* | Cattle A, E, G, J;<br>Sheep A, E | Spring 2019 (Apr, May), Autumn 2019 (Dec), Spring 2020 (Jun), Winter 2020 (Jan, Feb) | A,E | feces (6), feed (1), bedding (4), food trough (2) |
|  | 4 | *dfrD,lnuA,tetM* | Sheep E | Autumn 2019 (Dec), Winter 2020 (Mar), Spring 2020 (Jun) | A,E | feces (1), bedding (2), food trough (1) |
|  | 1 | *dfrD,lnuA,tetS,tetM* | Goat D | Winter 2020 (Mar) | A | feces (1) |
|  | 1 | *lnuA,tetM* | Cattle J | Autumn 2019 (Dec) | E | feed (1) |
| *L. aquatica* | 1 | *dfrK* | Sheep A | Spring 2019 (May) | A | feces (1) |

<sup>1</sup>gene traits of antibiotic resistance towards: folate inhibitors (*dfrD*, *dfrK*); lincosamides (*lnuA*); macrolides (*ermG,mefA,msrD*); tetracyclines (*tetM*, *tetS*)

### Impact of farm type, season, and farming practices in the prevalence of *L. monocytogenes*

*L. monocytogenes* was detected more frequently in cattle (7/10 positive farms, prevalence 4.1%) than in sheep (2/5 positive farms, prevalence 4.5%) and goat farms (1/4 positive farms, prevalence 0.2%) (Table S2). Although *L. monocytogenes* was detected more frequently in “Cattle G” and “Sheep B” farms, no remarkable differences in management practices were detected compared to other farms of the same species where the prevalence of *Listeria* spp. was lower (Table S1). Since only 1 out of 4 goat farms was positive for *L. monocytogenes* (Fig. 1C), and the prevalence was extremely low in this farm (0.7%), goat farms were not included in further statistical analyses.

*L. monocytogenes* presence in consecutive seasons was only detected in farm “Cattle G” (Fig. 1A). In cattle farms the overall prevalence was higher in winter than in autumn (*P*<0.05) (Fig. 1B and Fig. 4A). In sheep farms the overall prevalence was higher in winter and spring than in autumn (*P*<0.05) (Fig. 1B and Fig. 4B). Interestingly, cows were more likely to shed *L. monocytogenes* on the second lactation than on the first, forth or higher lactations (*P*<0.05) (Fig. 4C), but DIM did not impact the frequency of *L. monocytogenes* fecal shedding (Fig. 4D). In sheep, no significant association was found between the lactation number and frequency of *L. monocytogenes* fecal shedding (Fig. 4E). Differences in production hygiene were observed between both cattle and sheep farms, but there was no significant correlation between hygiene scores and *L. monocytogenes* prevalence (Table S5).

**Figure 4.**
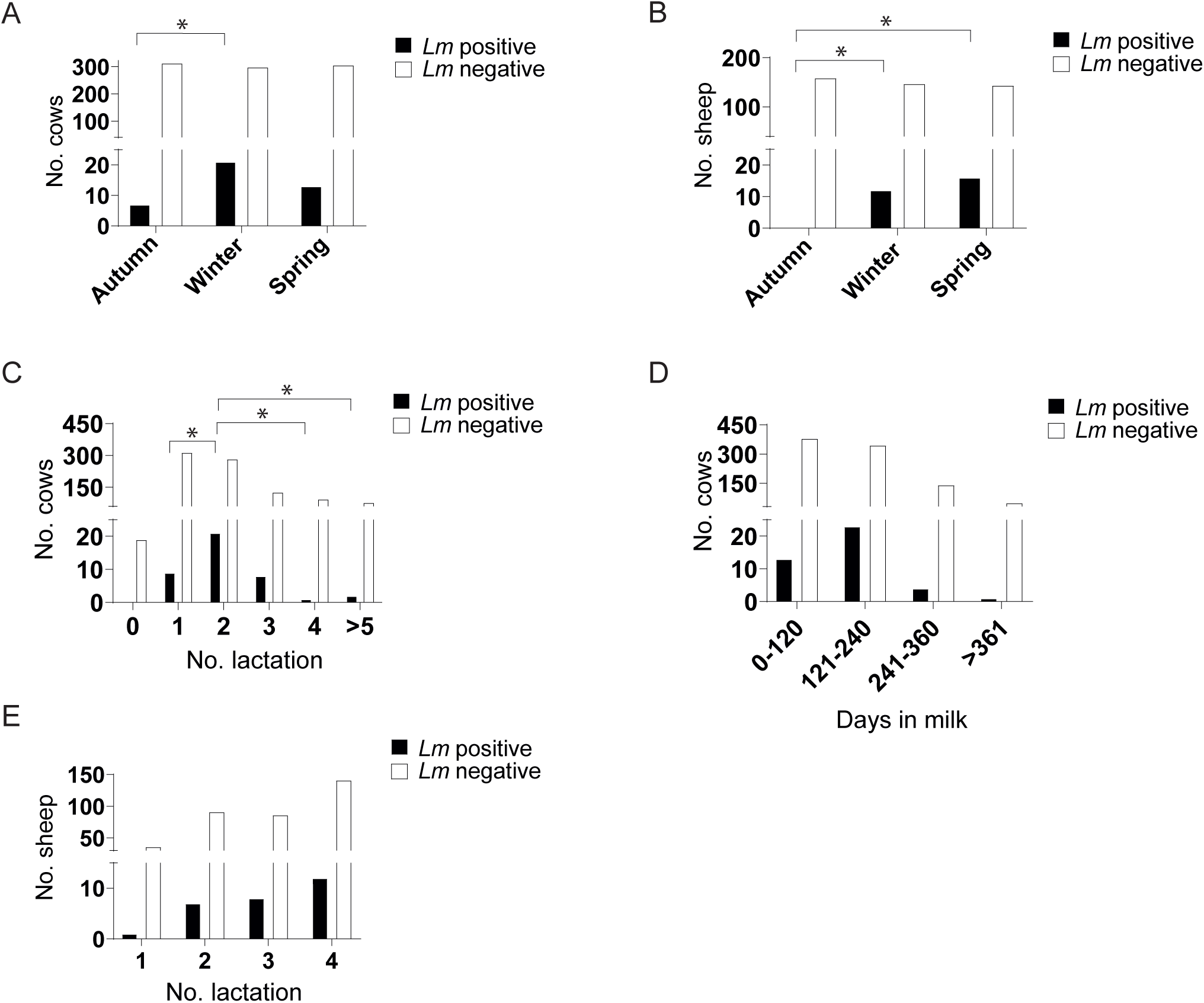
Impact of seasons, lactation number and days in milk (DIM) in the number of *L. monocytogenes* fecal shedders. A, B) Number of *L. monocytogenes* fecal shedders per season in cattle and sheep farms, respectively. C, E) Number of *L. monocytogenes* fecal shedders per lactation in cattle and sheep farms, respectively. D) Number of *L. monocytogenes* fecal shedders per DIM range in cattle farms. Statistically significant differences were evaluated by Chi-square tests. Stars denote P-values < 0.05.

Genotypes isolated from feces samples did not occur in MFS, indicating that pre-milking teat disinfection used in these farms was effective to prevent *L*. *monocytogenes* milk contamination, with exception for farm “Sheep B”, in which this procedure was not typically performed (similarly to all small ruminants’ dairy farms). Accordingly, L2-SL7-ST7-CT5782 isolated from sheep feces samples was identified in MFS in the farm “Sheep B”.

Although all farms reported usage of antibiotics for treatment purposes (Table S1), acquired genetic traits of antibiotic resistance were rare in *L. monocytogenes.* Interestingly, among *L. innocua,* 19 isolates harbored *tetM* genes, 13 (68.4%) of which were isolated from farms using tetracyclines.

## DISCUSSION

Understanding *L. monocytogenes* population dynamics and its biodiversity is essential for effective disease surveillance and the development of control strategies. To the best of our knowledge, this is the largest longitudinal study on the prevalence, ecology and genomic characteristics of *L. monocytogenes* in individual dairy ruminants and the farm environment. Other reports have either applied a longitudinal study design with a reduced number of farms (one to three farms) (7, 9, 10, 36) and/or analyzed feces samples from randomly chosen farm ruminants, which limits the understanding of global and individual fecal shedding patterns over time (9, 10).

In this study, the prevalence of *L. monocytogenes* detected in dairy farms (3.8% in feces samples and 2.5% in farm environment samples) was lower than previously reported in dairy farms with no clinical listeriosis cases (fecal sample prevalence of 0-60% in cattle and 14.2% in sheep) in farms from USA and Europe (6–9, 34, 36). Differences in climate and farm management (e.g., feed used) between different geographical regions may account for the low prevalence of *L. monocytogenes* in our study compared to previous studies performed in northern countries (37–39).

*L. monocytogenes* was detected more frequently in cattle farms than in small-ruminant farms, which is coherent with previous studies regarding the epidemiology of listeriosis among cattle and small ruminants (8, 11, 34, 40). Interestingly, the pathogenic species *L. ivanovii*, reported in small ruminants in previous studies (40–42) was not detected in any of our farms, which could be due to its relative low prevalence (42, 43) or to possible biases of isolation protocols that have typically been optimized for recovery of *L. monocytogenes* (42). Our results are in line with reports showing that the incidence of *L. innocua* in ruminants feces is higher (9.7-22.7%) than that of *L. monocytogenes* (1.8-9.3%) (17, 44). Of note, it has been previously shown that competitive *L. innocua* strains could mask detection of *L. monocytogenes* during enrichment protocols (45).

Although consumption of spoiled silage is thought to be the principal source of infection for ruminants (3), there is no evident link between listeriosis and silage feeding in up to a third of animal listeriosis cases (46). In this study, in 50% of the farms where fecal shedders were detected, no *L. monocytogenes* could be detected in feed, food troughs or water troughs. It has been suggested that contaminated water or feed by wildlife, birds, insects, farm staff, visitors, farm transport of animals, or farm equipment could vehiculate *L. monocytogenes* to farms (10, 13, 34, 47, 48). Moreover, introduction of new animals could be a potential source for the introduction of new *Listeria* strains into the farm, which can be transmitted via fecal-oral contact to other animals of the herd.

The majority of isolates retrieved here belonged to lineage I (particularly to SL1/CC1, SL219/CC4 and SL87/CC87) which is significantly associated with a clinical origin both in humans and animals (29, 49, 50). CC1 has been shown to be highly associated with dairy products (30, 51) and cause human listeriosis outbreaks (52, 53). CC1 is hypervirulent and colonizes better the intestinal lumen and invades more intestinal tissues than hypovirulent clones (CC9 and CC121) (30, 31, 50). CC87 has been previously reported as predominant in foodborne and clinical isolates in China and related to two outbreaks in Northern Spain (54–56).

Previously, *L. monocytogenes* isolates of identical or nearly identical genotypes (based on “multivirulence” locus sequence typing or pulsed-field gel electrophoresis) have been described to occur for years on dairy farms, although the majority of the genotypes were sporadic (9, 10, 57). In the present study, by using a genome-based approach, the same genotypes were found in multiple animals and surfaces within the same farms. Interestingly, with exception for one sheep, identical genotypes could not be detected in the same animal along different seasons, suggesting that fecal shedding period is shorter than the time frame between samplings dates. Indeed, previous experimental studies have shown that fecal shedding lasted 10 days in sheep inoculated orally with a high dose of *L. monocytogenes* (10^10^ colony forming units) (58). Accordingly, fecal carriage of *L. monocytogenes* in healthy human adults is reported to be transient (59). However, studies in wild and domestic ruminants suggest that animals can silently carry *L. monocytogenes* in tonsils even without fecal shedding (58, 60). We hypothesize that these silent carriers could under certain circumstances (i.e. immunosuppression, stress factors) shed this bacterium from the tonsils to the environment through feces leading to intermittent shedding. Tonsil infection may also be a portal of entry for infection in animals and may explain why *L. monocytogenes* was not identified in the feces of sheep herd where a listeriosis outbreak occurred.

Although *L. monocytogenes* could be isolated from cattle all year round, the overall prevalence of *L. monocytogenes* in cattle farms was significantly higher in samples collected during winter which is in line with previous reports (9, 61, 62). Cows in their second lactation had a higher probability of *L. monocytogenes* fecal shedding than cows on the first, forth or higher lactations. An inadequate transition from the first to the second lactation with a poor body condition score could impair immune function (63) and predispose to *L. monocytogenes* colonization. Interestingly, *tetM* genes were detected more frequently in *L. innocua* isolates from farms using tetracycline than in farms not using this antibiotic, highlighting the importance of antibiotic stewardship in veterinary medicine.

In summary, our data show that i) *L. innocua* and *L. monocytogenes* were the most prevalent *Listeria* spp. in both dairy ruminant feces and farm-associated environments; *(ii)* single ruminants can harbor *L. monocytogenes* alone or together with *L. innocua* without clinical signs of infection; *(iii) L. monocytogenes* could be isolated from half of the dairy farms sampled; *(iv) L. monocytogenes* fecal shedding can present high levels of month-to-month variation and could occur as an isolated sporadic case or as a high-number of fecal shedders case; *(v)* except one isolated case, there was no evidence of transmission of *L. monocytogenes* CTs between farms, indicating intra-farm transmission dynamics; (*vi*) CC1 and CC4 hypervirulent *L. monocytogenes* clones, which are among the most common *L. monocytogenes* CCs responsible for human infection, represented 30% of the *L. monocytogenes* isolates retrieved in this study and were mainly obtained from host associated samples (feces); (*vii*) the overall *L. monocytogenes* prevalence was higher in winter than in autumn in cattle farms and higher in winter and spring than autumn in sheep farms; and (*viii*) cows were more likely to shed *L. monocytogenes* on the second lactation than on the first, forth or higher lactations.

Our data are consistent with the hypothesis that dairy farms may be sites that favor the selection of invasive *L. monocytogenes* clones, which are indeed shed in the feces more efficiently than hypovirulent clones (30), and constitute a reservoir for hypervirulent strains that can colonize dairy products. This study improves the understanding of *Listeria* spp. prevalence and ecology in the dairy ruminant environment and may contribute to the development of effective disease surveillance and control strategies to reduce the number of both human and animal listeriosis cases.

## MATERIALS AND METHODS

### Farms

The study population consisted of 19 dairy ruminant farms (10 cattle, 5 sheep, and 4 goat) with different housing systems, management practices, and herd sizes located in the provinces (administrative division in Spain) of Valencia, Alicante, Castellón, Murcia, and Albacete (mid-east and south-east of Spain) (Table S1).

No history of clinical listeriosis had been observed in any of the farms before and/or during the sampling period, except for farm “Sheep C” which suffered a listeriosis outbreak in the last season sampled (spring 2020).

### Sample collection

During the sampling period (winter 2018 to spring 2019 and winter 2019 to spring 2020), each farm was visited once per season (autumn, winter, and spring), for a total of 3 visits per farm. Farm characteristics, sampling dates, and *Listeria* spp. isolated are indicated on Tables S1 and S3. Farm “Sheep B” was sampled 11 times during 7 consecutive seasons from autumn 2018 to spring 2020 for *Listeria* genomic diversity analysis due to the discovery of the new species *L. valentina* in the February/2019 sampling (1). For consistency with data from other farms, only data from three consecutive seasons (autumn 2018-Nov-07, winter 2019-Feb-27, and spring 2019-Apr-10) in this farm were considered for prevalence and statistics calculations.

On each farm visit, 50 samples (32 samples of feces from individual animals, 3 samples of feed, 3 samples of bedding, 3 MFS, and 9 surface swabs (3 from milking station floor, 3 from water troughs, 3 from food troughs) were collected during three consecutive seasons (autumn, winter, and spring) amounting to 150 samples per farm. The same 32 animals sampled during the first farm visit, were monitored in the course of followings evaluations (three seasons total) by veterinarians during the usual handling of the animals, following the guidelines of European Union Directive 2010/63/EU for the protection of animals used for scientific purposes (64). Cows, sheep, or goats that were sold in the intervals between the sampling periods were replaced in the study with a new animal.

MFS were selected for sampling since the prevalence of *L. monocytogenes* is twice that in BTM(9). Each sample was collected into a sterile bag by the use of clean gloves or sampling utensils. Rectal fecal grab samples were collected from randomly selected animals in each farm to have a representation of all lactation numbers. Fecal samples were obtained by rectal grab to avoid cross-contamination among animals. This routine veterinary practice does not require the approval of the Animal Ethics and Experimentation Committee. Bedding samples, food troughs samples, water troughs samples, and milking station floor samples were collected from diverse locations on each farm. All samples were collected using disposable gloves by aseptic conditions and stored in clean coolers with ice packs for transit to the laboratory. Samples were processed within 2-12 h of collection.

### Animal cleanliness and production hygiene

A numerical scoring system for assessing animal cleanliness of 5 body areas (tail head, ventral abdomen, udder, upper rear limb, and lower rear limb) was used for the individual animals as previously described (scale of 1 to 5, where score 1 = very clean, score 5 = heavily soiled) (65). Production hygiene was evaluated based on the cleanliness of the premises (milk room, milking station, feed troughs, water troughs and beddings) on farm visits on a scale of 1 to 3 as previously described (9). A score of “1” corresponded to a major deficit in production hygiene, “2” to a minor deficit in production hygiene, and “3” to no notable deficit in production hygiene.

### *Listeria* spp. isolation and identification

*Listeria* spp. were isolated as previously described (1, 60). Briefly, 8 g of rectal fecal samples or bedding samples were diluted 1/10 in Half-Fraser broth (Scharlab, Spain), homogenized and incubated at 30 °C for 24 h for enrichment. Swab samples (feed troughs, water troughs, and milking station floor) were placed in 10 ml Half Fraser broth, vortexed for 2 min, and incubated at 30 °C for 24 h for enrichment. Entire MFS socks or 8 g of feed samples were used as sample material for primary enrichment in Half Fraser broth (30 °C, 24h).

Samples were homogenized manually for 1 min until the solid matter was completely suspended in the enrichment solution. One hundred microlitres of the incubated suspension were transferred to 10 ml Fraser broth (Scharlab, Spain) and incubated at 37 °C for 24 h. After the second enrichment, 100 μl enriched culture and two tenfold dilutions were transferred to RAPID’*L.mono* plates (BioRad, USA) and incubated at 37 °C for 24 h. Characteristic *Listeria* spp. colonies (colonies were blue or white, with or without a yellow halo, round, convex, 1 to 2 mm) were confirmed in selective Oxford agar plates for *Listeria* (Scharlab, Spain) (colonies were approximately 2 mm in diameter, grey-green with a black sunken center and a black halo) and Columbia CNA agar with 5% sheep blood agar plates (colonies were opaque, flat, 1 to 2 mm). From each positive sample, one isolate colony was obtained and preserved in glycerol at - 80°C and sent to the World Health Organization Collaborating Centre (WHOCC) for *Listeria*, Institut Pasteur, Paris, France, for species identification and characterization. *Listeria* isolates were identified with matrix-assisted laser desorption ionization-time of flight mass spectrometry using the MicroFlex LT system with the last MBT library DB-7854 (Bruker Daltonics, Germany), as previously described (66) and by whole genome sequencing as previously described (1).

### Genome sequencing and assembly

DNA extraction was carried out with the NucleoSpin Tissue purification kit (Macherey-Nagel, Germany) from 0.9 mL Brain heart infusion (Difco, USA) cultures grown overnight at 35°C. DNA libraries were prepared using the Nextera XT DNA Sample kit (Illumina, USA), and sequenced in a NextSeq 500 platform (Illumina, USA) using 2 x 150 bp runs, according to the manufacturer’s protocol. Raw reads were trimmed with fqCleaner v.3.0 (Alexis Criscuolo, Institut Pasteur, Paris) as previously described (1, 60), and assembled with SPAdes v.3.12.0 (67) using automatic k-mer selection and the --only-assembler and --careful options.

### Molecular typing and phylogenetic analysis

*In silico* typing was performed from the assemblies using the genoserogrouping (26), MLST (7 loci (27)), cgMLST profiles (1,748 loci (28)), resistance and virulence schemes (244 loci) implemented at using BIGSdb-*L. monocytogenes* v.1.30 (https://bigsdb.pasteur.fr/listeria; (28, 68)). Genes were scanned at BIGSdb-*L. monocytogenes* using the BLASTN algorithm, with minimum nucleotide identity and alignment length coverage of 70% and word size of 10, as previously described (28). MLST profiles were classified into ST and grouped into CCs as previously described (27). cgMLST profiles were grouped into CTs and SLs, using the cut-offs of 7 and 150 allelic mismatches, respectively, as previously described (28). Minimum spanning trees and single linkage dendrograms were built from cgMLST profiles using Bionumerics 7.6 software (Applied Maths, Belgium) and annotated with iTol v.4.2 (69). Assemblies were also screened for antimicrobial resistance genes and the presence of plasmids using ABRicate v.1.0.1 (70–74) and MOB-suite v.2.0.1 (75), respectively.

### Statistical analysis

For DIM analysis cows were grouped into different categories considering the dairy cattle lactation curve and classified in Early lactation (0 -120 days), Mid-lactation (121 – 240 days), Late lactation (241 – 360) and End of Lactation (> 361 DIM). DIM were not analyzed in sheep farms since all the ewes in the same farm were synchronized using intravaginal sponges and delivered approximately the same day. Shannon diversity indices and Hutcheson T-test were calculated using the web https://www.dataanalytics.org.uk/comparing-diversity/ (76). The rest of statistical analyses were conducted with IBM SPSS Statistics version 25. The significance level for all statistical tests was *P*<0.05. Chi-square tests were performed to determine the effect of season, the number of lactation and DIM on the number of *L. monocytogenes* fecal shedders. Spearman’s rank-order correlations were done to evaluate the association between the farm hygiene score and *L. monocytogenes* prevalence.

## Data availability

Sequences obtained in this study will be made publicly available at the European Nucleotide Archive (BioProjects PRJEB45781 and PRJEB36008) and BIGSdb-*Listeria* (bigsdb.pasteur.fr/listeria) upon publication. The accession numbers are detailed in Table S3.

## Acknowledgments

We are grateful for the willingness of the farmers to participate in this study. This work was supported by Generalitat Valenciana (Project reference GV/2018/A/183), the Spanish Ministry of Science and Innovation (Project reference PID2019-110764RA-I00/AEI/10.13039/501100011033) and Universidad CEU Cardenal Herrera Programa INDI 20/40 (JJQ), the Spanish Ministry of Science and Innovation (Project reference PID2020-119462RA-I00/AEI/10.13039/ 501100011033) (AGM), Institut Pasteur, Inserm, and Santé Publique France (ML). J.J. Quereda is supported by a “Ramón y Cajal” contract of the Spanish Ministry of Science, Innovation, and Universities (RYC-2018-024985-I). C. Palacios-Gorba is supported by a Predoctoral contract from the Universidad Cardenal Herrera-CEU. The funders had no role in study design, data collection, and interpretation, or the decision to submit the work for publication.

---

**Table S1.** Characteristics and management practices of the investigated dairy farms.

| Farm | Province | Herd size | Area per animal (m<sup>2</sup>) | Silage | Bedding material | Sampling periods | Antibiotics used |
| --- | --- | --- | --- | --- | --- | --- | --- |
| Cattle A | Valencia | >400 | 17 | Corn+Barley | Compost | 2018-10-17 / 2019-02-11 / 2019-04-08 | Penicillin, ceftiofur, trimethoprim, sulfonamide, tylosin, marbofloxacin |
| Cattle B | Valencia | >400 | 15 | Corn | Compost | 2018-11-14 / 2019-02-19 / 2019-05-27 | Ceftiofur, penicillin, oxitetracycline |
| Cattle C | Murcia | >400 | 23 | Corn | Compost | 2018-11-15 / 2019-03-20 / 2019-05-13 | Ceftiofur, marbofloxacin, amoxicillin |
| Cattle D | Albacete | 40 | 8.1 | Corn | Straw | 2018-12-17 / 2019-02-08 / 2019-03-26 | Ceftiofur, florfenicol |
| Cattle E | Valencia | 124 | 8 | No | Straw | 2018-10-23 / 2019-02-27 / 2019-04-30 | Ceftiofur, marbofloxacin, trimethoprim, sulfonamide, penicillin, dihydrostreptomycin, enrofloxacin |
| Cattle F | Albacete | 190 | 263 | Oats | Straw | 2019-10-28 / 2020-02-21 / 2020-06-17 | Florfenicol, trimethoprim, sulfonamide, penicillin |
| Cattle G | Murcia | >400 | 22 | Oats | Compost | 2019-10-28 / 2020-02-21 / 2020-06-17 | Sulfonamide, marbofloxacin, tetracycline, enrofloxacin |
| Cattle H | Albacete | >400 | 3.3 | Corn+Oats | Compost | 2019-11-19 / 2020-02-28 / 2020-05-20 | Trimethoprim, sulfonamide, penicillin, tylosin, oxitetracycline |
| Cattle I | Valencia | 150 | 3.3 | Corn | Mattress+wood-shavings | 2019-12-10 / 2020-02-27 / 2020-05-26 | Florfenicol, trimethoprim, sulfonamide, penicillin, tylosin, oxitetracycline |
| Cattle J | Valencia | >400 | 22.5 | No | Straw | 2019-12-16 / 2020-01-27 / 2020-05-26 | Oxitetracycline |
| Sheep A | Albacete | 1400 | 1.8 | No | Straw | 2018-11-23 / 2019-01-17 / 2019-05-06 | Amoxicillin |
| Sheep B | Castellon | >2000 | 2 | Alfalfa | Straw | 2018-11-07 / 2019-02-27 / 2019-04-10 | Amoxicillin, oxitetracycline, marbofloxacin, trimethoprim, sulfonamide, tulathromycin |
| Sheep C | Albacete | >2000 | 4 | Corn | Straw | 2019-11-05 / 2020-01-13 / 2020-05-15 | Amoxicillin, oxytetracycline |
| Sheep D | Albacete | >2000 | 3 | Corn | Straw | 2019-12-03 / 2020-02-17 / 2020-05-20 | Amoxicillin |
| Sheep E | Alicante | >2000 | 3 | Wet brewery grains | Straw | 2019-12-17 / 2020-03-09 / 2020-06-17 | Cephalosporin, oxitetracycline, colistin |
| Goat A | Castellón | 400 | 1.06 | No | Straw | 2018-11-01 / 2019-01-23 / 2019-04-16 | Enrofloxacin, amoxicillin |
| Goat B | Murcia | 600 | 3 | No | Straw | 2018-12-10 / 2019-04-01 / 2019-05-10 | Enrofloxacin, oxitetracycline, tylosin, erythromycin |
| Goat C | Albacete | 750 | 1.64 | Corn | Straw | 2019-11-05 / 2020-01-27 / 2020-05-20 | Terramycin |
| Goat D | Murcia | >2000 | 4 | Corn | Straw | 2019-11-26 / 2020-03-13 / 2020-06-19 | Amoxicillin, gentamycin, ceftiofur, enrofloxacin |

All farms had an open-air barn structure, a parlor milking system, and used a conventional farming system except farm “Cattle F” that followed an organic farming system. All farms used a loose housing system (bedded-pack), except farms “Cattle H” and “Cattle I” which used a free stall system. All farms used a mechanic scrape as a feces removal system.

**Table S2.**
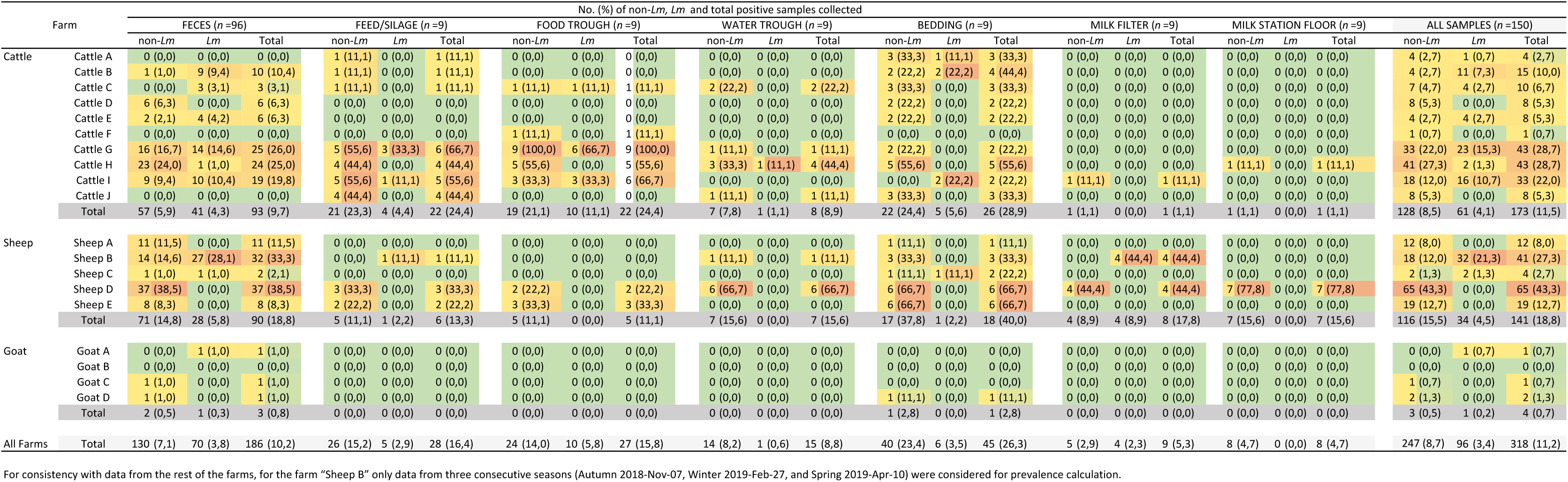
Prevalence of non-pathogenic *Listeria*, *L. monocytogenes* (*Lm* ), and total *Listeria* spp. in feces samples and the farm environment from 19 Spanish dairy farms.

| Farm |  | No. (%) of non-*Lm*, *Lm* and total positive samples collected |  |  |  |  |  |  |  |  |  |  |  |  |  |  |  |  |  |  |  |  |  |  |  |
| --- | --- | --- | --- | --- | --- | --- | --- | --- | --- | --- | --- | --- | --- | --- | --- | --- | --- | --- | --- | --- | --- | --- | --- | --- | --- |
|  |  | FECES (n =96) |  |  | FEED/SILAGE (n =9) |  |  | FOOD TROUGH (n =9) |  |  | WATER TROUGH (n =9) |  |  | BEDDING (n =9) |  |  | MILK FILTER (n =9) |  |  | MILK STATION FLOOR (n =9) |  |  | ALL SAMPLES (n =150) |  |  |
|  |  | non-*Lm* | *Lm* | Total | non-*Lm* | *Lm* | Total | non-*Lm* | *Lm* | Total | non-*Lm* | *Lm* | Total | non-*Lm* | *Lm* | Total | non-*Lm* | *Lm* | Total | non-*Lm* | *Lm* | Total | non-*Lm* | *Lm* | Total |
| Cattle | Cattle A | 0 (0,0) | 0 (0,0) | 0 (0,0) | 1 (11,1) | 0 (0,0) | 1 (11,1) | 0 (0,0) | 0 (0,0) | 0 (0,0) | 0 (0,0) | 0 (0,0) | 0 (0,0) | 3 (33,3) | 1 (11,1) | 3 (33,3) | 0 (0,0) | 0 (0,0) | 0 (0,0) | 0 (0,0) | 0 (0,0) | 0 (0,0) | 4 (2,7) | 1 (0,7) | 4 (2,7) |
|  | Cattle B | 1 (1,0) | 9 (9,4) | 10 (10,4) | 1 (11,1) | 0 (0,0) | 1 (11,1) | 0 (0,0) | 0 (0,0) | 0 (0,0) | 0 (0,0) | 0 (0,0) | 0 (0,0) | 2 (22,2) | 2 (22,2) | 4 (44,4) | 0 (0,0) | 0 (0,0) | 0 (0,0) | 0 (0,0) | 0 (0,0) | 0 (0,0) | 4 (2,7) | 11 (7,3) | 15 (10,0) |
|  | Cattle C | 0 (0,0) | 3 (3,1) | 3 (3,1) | 1 (11,1) | 0 (0,0) | 1 (11,1) | 1 (11,1) | 1 (11,1) | 1 (11,1) | 2 (22,2) | 0 (0,0) | 2 (22,2) | 3 (33,3) | 0 (0,0) | 3 (33,3) | 0 (0,0) | 0 (0,0) | 0 (0,0) | 0 (0,0) | 0 (0,0) | 0 (0,0) | 7 (4,7) | 4 (2,7) | 10 (6,7) |
|  | Cattle D | 6 (6,3) | 0 (0,0) | 6 (6,3) | 0 (0,0) | 0 (0,0) | 0 (0,0) | 0 (0,0) | 0 (0,0) | 0 (0,0) | 0 (0,0) | 0 (0,0) | 0 (0,0) | 2 (22,2) | 0 (0,0) | 2 (22,2) | 0 (0,0) | 0 (0,0) | 0 (0,0) | 0 (0,0) | 0 (0,0) | 0 (0,0) | 8 (5,3) | 0 (0,0) | 8 (5,3) |
|  | Cattle E | 2 (2,1) | 4 (4,2) | 6 (6,3) | 0 (0,0) | 0 (0,0) | 0 (0,0) | 0 (0,0) | 0 (0,0) | 0 (0,0) | 0 (0,0) | 0 (0,0) | 0 (0,0) | 2 (22,2) | 0 (0,0) | 2 (22,2) | 0 (0,0) | 0 (0,0) | 0 (0,0) | 0 (0,0) | 0 (0,0) | 0 (0,0) | 4 (2,7) | 4 (2,7) | 8 (5,3) |
|  | Cattle F | 0 (0,0) | 0 (0,0) | 0 (0,0) | 0 (0,0) | 0 (0,0) | 0 (0,0) | 1 (11,1) | 0 (0,0) | 1 (11,1) | 0 (0,0) | 0 (0,0) | 0 (0,0) | 0 (0,0) | 0 (0,0) | 0 (0,0) | 0 (0,0) | 0 (0,0) | 0 (0,0) | 0 (0,0) | 0 (0,0) | 0 (0,0) | 1 (0,7) | 0 (0,0) | 1 (0,7) |
|  | Cattle G | 16 (16,7) | 14 (14,6) | 25 (26,0) | 5 (55,6) | 3 (33,3) | 6 (66,7) | 9 (100,0) | 6 (66,7) | 9 (100,0) | 1 (11,1) | 0 (0,0) | 1 (11,1) | 2 (22,2) | 0 (0,0) | 2 (22,2) | 0 (0,0) | 0 (0,0) | 0 (0,0) | 0 (0,0) | 0 (0,0) | 0 (0,0) | 33 (22,0) | 23 (15,3) | 43 (28,7) |
|  | Cattle H | 23 (24,0) | 1 (1,0) | 24 (25,0) | 4 (44,4) | 0 (0,0) | 4 (44,4) | 5 (55,6) | 0 (0,0) | 5 (55,6) | 3 (33,3) | 1 (11,1) | 4 (44,4) | 5 (55,6) | 0 (0,0) | 5 (55,6) | 0 (0,0) | 0 (0,0) | 0 (0,0) | 1 (11,1) | 0 (0,0) | 1 (11,1) | 41 (27,3) | 2 (1,3) | 43 (28,7) |
|  | Cattle I | 9 (9,4) | 10 (10,4) | 19 (19,8) | 5 (55,6) | 1 (11,1) | 5 (55,6) | 3 (33,3) | 3 (33,3) | 6 (66,7) | 0 (0,0) | 0 (0,0) | 0 (0,0) | 0 (0,0) | 2 (22,2) | 2 (22,2) | 1 (11,1) | 0 (0,0) | 1 (11,1) | 0 (0,0) | 0 (0,0) | 0 (0,0) | 18 (12,0) | 16 (10,7) | 33 (22,0) |
|  | Cattle J | 0 (0,0) | 0 (0,0) | 0 (0,0) | 4 (44,4) | 0 (0,0) | 4 (44,4) | 0 (0,0) | 0 (0,0) | 0 (0,0) | 1 (11,1) | 0 (0,0) | 1 (11,1) | 3 (33,3) | 0 (0,0) | 3 (33,3) | 0 (0,0) | 0 (0,0) | 0 (0,0) | 0 (0,0) | 0 (0,0) | 0 (0,0) | 8 (5,3) | 0 (0,0) | 8 (5,3) |
|  | Total | 57 (5,9) | 41 (4,3) | 93 (9,7) | 21 (23,3) | 4 (4,4) | 22 (24,4) | 19 (21,1) | 10 (11,1) | 22 (24,4) | 7 (7,8) | 1 (1,1) | 8 (8,9) | 22 (24,4) | 5 (5,6) | 26 (28,9) | 1 (1,1) | 0 (0,0) | 1 (1,1) | 1 (1,1) | 0 (0,0) | 1 (1,1) | 128 (8,5) | 61 (4,1) | 173 (11,5) |
| Sheep | Sheep A | 11 (11,5) | 0 (0,0) | 11 (11,5) | 0 (0,0) | 0 (0,0) | 0 (0,0) | 0 (0,0) | 0 (0,0) | 0 (0,0) | 0 (0,0) | 0 (0,0) | 0 (0,0) | 1 (11,1) | 0 (0,0) | 1 (11,1) | 0 (0,0) | 0 (0,0) | 0 (0,0) | 0 (0,0) | 0 (0,0) | 0 (0,0) | 12 (8,0) | 0 (0,0) | 12 (8,0) |
|  | Sheep B | 14 (14,6) | 27 (28,1) | 32 (33,3) | 0 (0,0) | 1 (11,1) | 1 (11,1) | 0 (0,0) | 0 (0,0) | 0 (0,0) | 1 (11,1) | 0 (0,0) | 1 (11,1) | 3 (33,3) | 0 (0,0) | 3 (33,3) | 0 (0,0) | 4 (44,4) | 4 (44,4) | 0 (0,0) | 0 (0,0) | 0 (0,0) | 18 (12,0) | 32 (21,3) | 41 (27,3) |
|  | Sheep C | 1 (1,0) | 1 (1,0) | 2 (2,1) | 0 (0,0) | 0 (0,0) | 0 (0,0) | 0 (0,0) | 0 (0,0) | 0 (0,0) | 0 (0,0) | 0 (0,0) | 0 (0,0) | 1 (11,1) | 1 (11,1) | 2 (22,2) | 0 (0,0) | 0 (0,0) | 0 (0,0) | 0 (0,0) | 0 (0,0) | 0 (0,0) | 2 (1,3) | 2 (1,3) | 4 (2,7) |
|  | Sheep D | 37 (38,5) | 0 (0,0) | 37 (38,5) | 3 (33,3) | 0 (0,0) | 3 (33,3) | 2 (22,2) | 0 (0,0) | 2 (22,2) | 6 (66,7) | 0 (0,0) | 6 (66,7) | 6 (66,7) | 0 (0,0) | 6 (66,7) | 4 (44,4) | 0 (0,0) | 4 (44,4) | 7 (77,8) | 0 (0,0) | 7 (77,8) | 65 (43,3) | 0 (0,0) | 65 (43,3) |
|  | Sheep E | 8 (8,3) | 0 (0,0) | 8 (8,3) | 2 (22,2) | 0 (0,0) | 2 (22,2) | 3 (33,3) | 0 (0,0) | 3 (33,3) | 0 (0,0) | 0 (0,0) | 0 (0,0) | 6 (66,7) | 0 (0,0) | 6 (66,7) | 0 (0,0) | 0 (0,0) | 0 (0,0) | 0 (0,0) | 0 (0,0) | 0 (0,0) | 19 (12,7) | 0 (0,0) | 19 (12,7) |
|  | Total | 71 (14,8) | 28 (5,8) | 90 (18,8) | 5 (11,1) | 1 (2,2) | 6 (13,3) | 5 (11,1) | 0 (0,0) | 5 (11,1) | 7 (15,6) | 0 (0,0) | 7 (15,6) | 17 (37,8) | 1 (2,2) | 18 (40,0) | 4 (8,9) | 4 (8,9) | 8 (17,8) | 7 (15,6) | 0 (0,0) | 7 (15,6) | 116 (15,5) | 34 (4,5) | 141 (18,8) |
| Goat | Goat A | 0 (0,0) | 1 (1,0) | 1 (1,0) | 0 (0,0) | 0 (0,0) | 0 (0,0) | 0 (0,0) | 0 (0,0) | 0 (0,0) | 0 (0,0) | 0 (0,0) | 0 (0,0) | 0 (0,0) | 0 (0,0) | 0 (0,0) | 0 (0,0) | 0 (0,0) | 0 (0,0) | 0 (0,0) | 0 (0,0) | 0 (0,0) | 0 (0,0) | 1 (0,7) | 1 (0,7) |
|  | Goat B | 0 (0,0) | 0 (0,0) | 0 (0,0) | 0 (0,0) | 0 (0,0) | 0 (0,0) | 0 (0,0) | 0 (0,0) | 0 (0,0) | 0 (0,0) | 0 (0,0) | 0 (0,0) | 0 (0,0) | 0 (0,0) | 0 (0,0) | 0 (0,0) | 0 (0,0) | 0 (0,0) | 0 (0,0) | 0 (0,0) | 0 (0,0) | 0 (0,0) | 0 (0,0) | 0 (0,0) |
|  | Goat C | 1 (1,0) | 0 (0,0) | 1 (1,0) | 0 (0,0) | 0 (0,0) | 0 (0,0) | 0 (0,0) | 0 (0,0) | 0 (0,0) | 0 (0,0) | 0 (0,0) | 0 (0,0) | 0 (0,0) | 0 (0,0) | 0 (0,0) | 0 (0,0) | 0 (0,0) | 0 (0,0) | 0 (0,0) | 0 (0,0) | 0 (0,0) | 1 (0,7) | 0 (0,0) | 1 (0,7) |
|  | Goat D | 1 (1,0) | 0 (0,0) | 1 (1,0) | 0 (0,0) | 0 (0,0) | 0 (0,0) | 0 (0,0) | 0 (0,0) | 0 (0,0) | 0 (0,0) | 0 (0,0) | 0 (0,0) | 1 (11,1) | 0 (0,0) | 1 (11,1) | 0 (0,0) | 0 (0,0) | 0 (0,0) | 0 (0,0) | 0 (0,0) | 0 (0,0) | 2 (1,3) | 0 (0,0) | 2 (1,3) |
|  | Total | 2 (0,5) | 1 (0,3) | 3 (0,8) | 0 (0,0) | 0 (0,0) | 0 (0,0) | 0 (0,0) | 0 (0,0) | 0 (0,0) | 0 (0,0) | 0 (0,0) | 0 (0,0) | 1 (2,8) | 0 (0,0) | 1 (2,8) | 0 (0,0) | 0 (0,0) | 0 (0,0) | 0 (0,0) | 0 (0,0) | 0 (0,0) | 3 (0,5) | 1 (0,2) | 4 (0,7) |
| All Farms | Total | 130 (7,1) | 70 (3,8) | 186 (10,2) | 26 (15,2) | 5 (2,9) | 28 (16,4) | 24 (14,0) | 10 (5,8) | 27 (15,8) | 14 (8,2) | 1 (0,6) | 15 (8,8) | 40 (23,4) | 6 (3,5) | 45 (26,3) | 5 (2,9) | 4 (2,3) | 9 (5,3) | 8 (4,7) | 0 (0,0) | 8 (4,7) | 247 (8,7) | 96 (3,4) | 318 (11,2) |

For consistency with data from the rest of the farms, for the farm “Sheep B” only data from three consecutive seasons (Autumn 2018-Nov-07, Winter 2019-Feb-27, and Spring 2019-Apr-10) were considered for prevalence calculation.

**Table S3.**
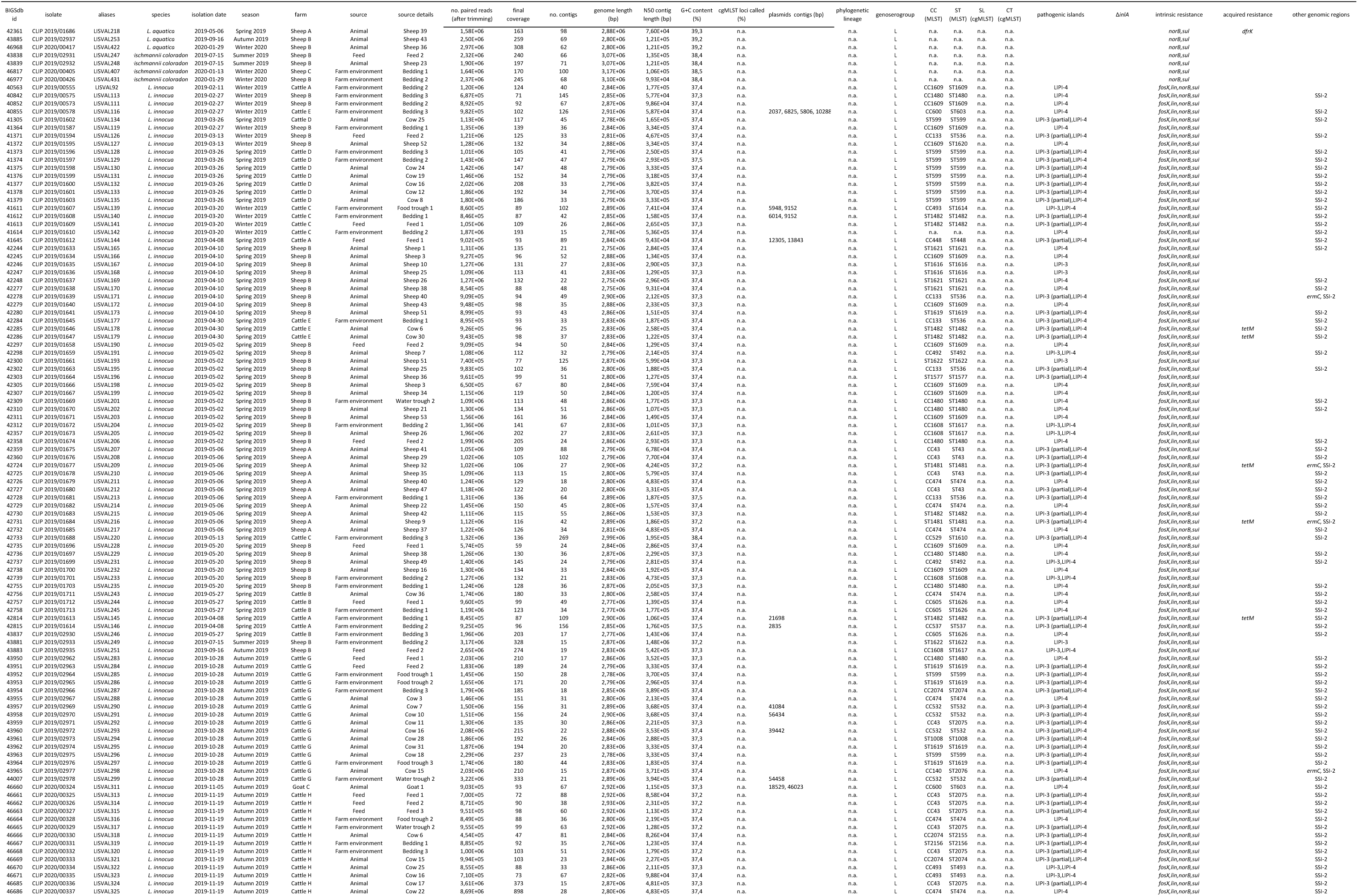
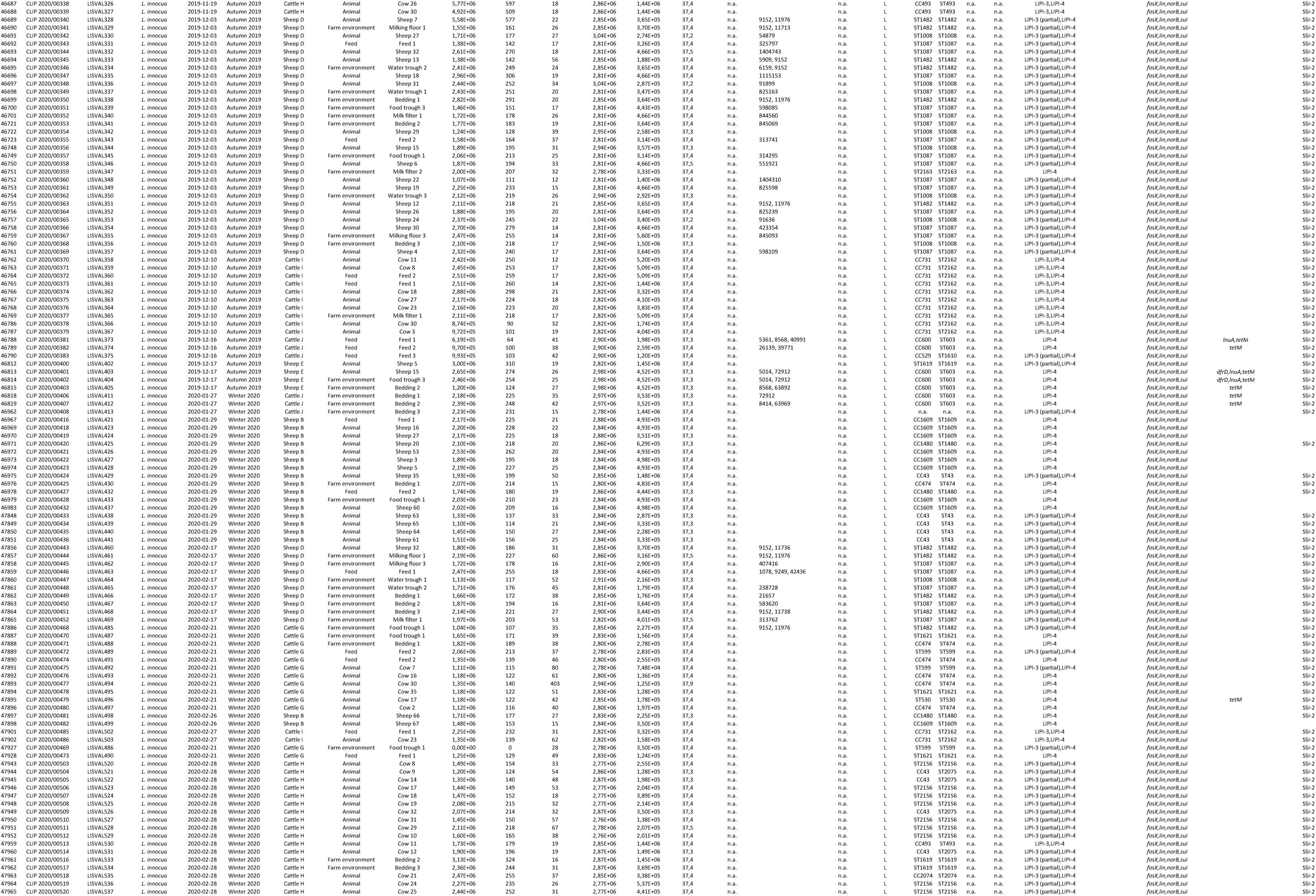
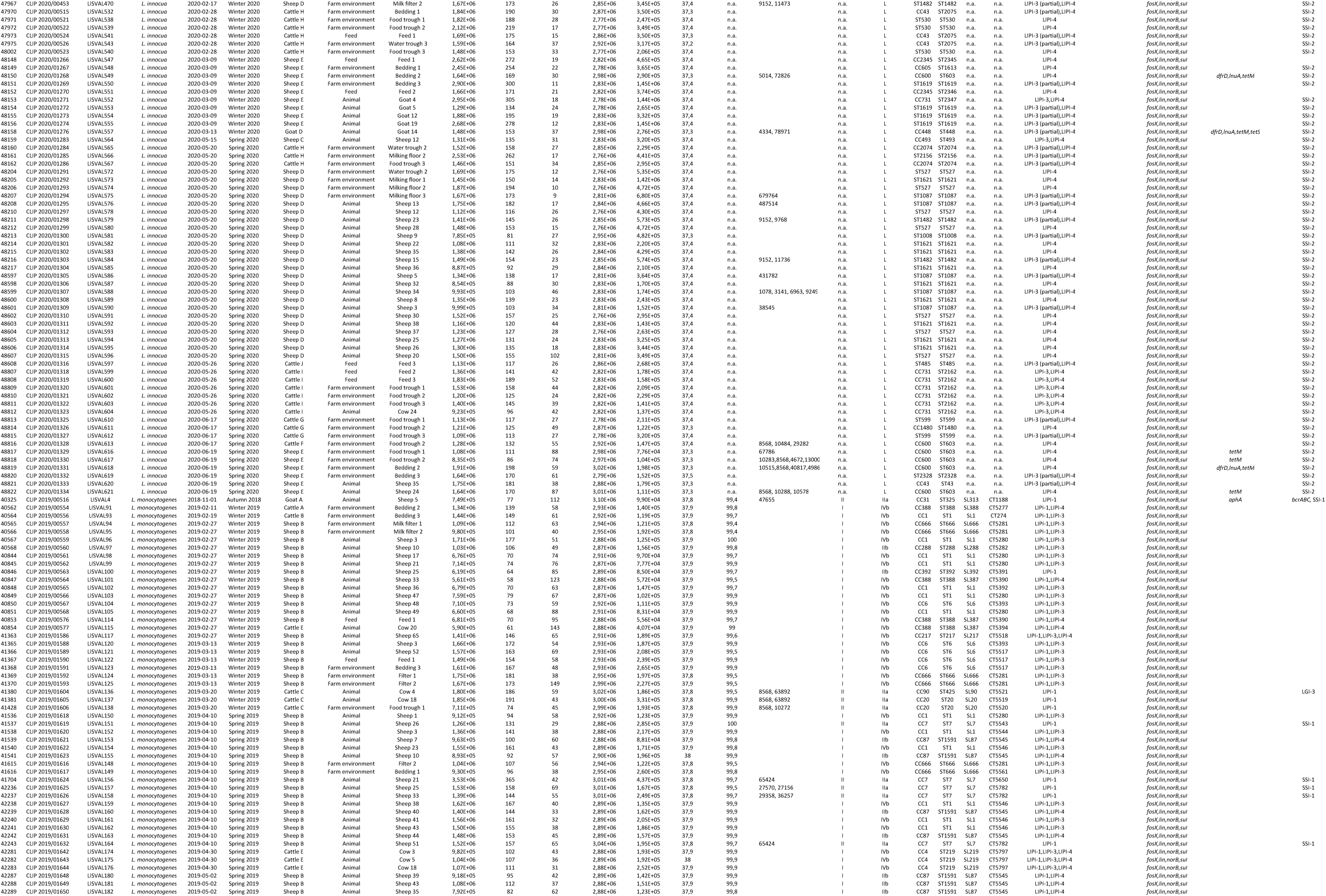
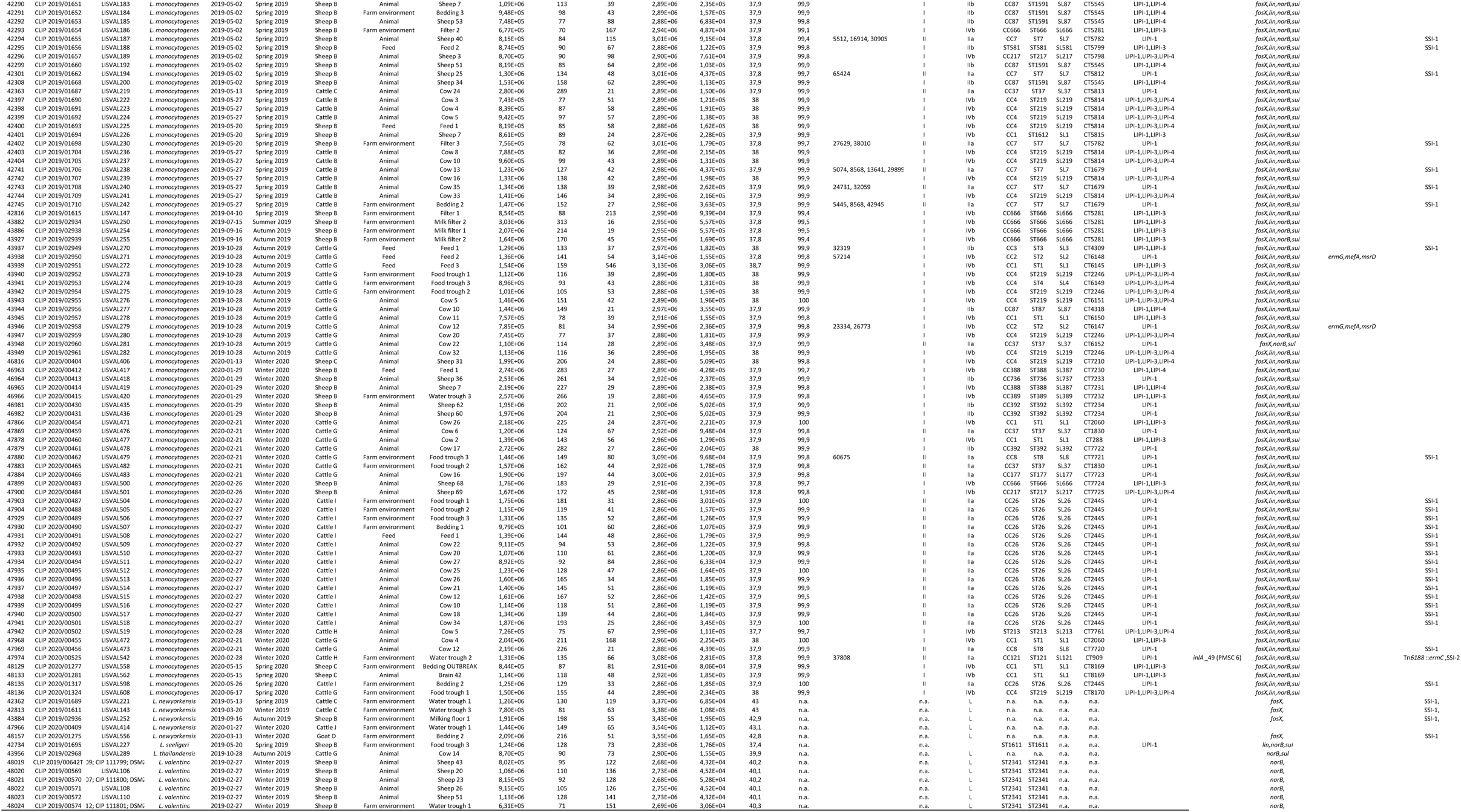
*Listeria* isolates characterized in this study.

**Table S4.**
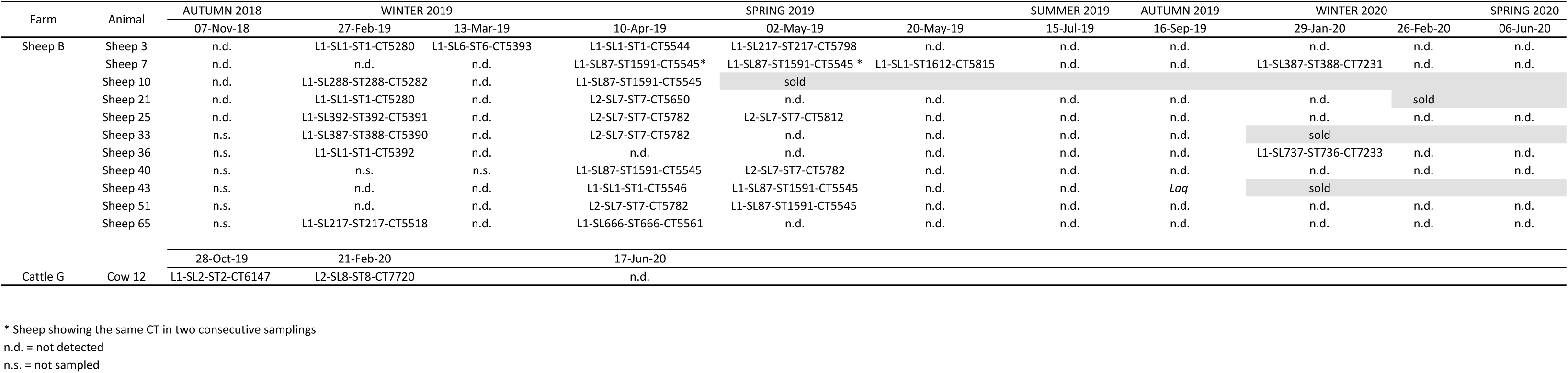
Individual animals where *Listeria monocytogenes* were isolated repeateadly in different samplings

**Table S5.** Correlation of hygienic scores and prevalence of *L. monocytogenes* on the investigated dairy farms. Hygienic scores were calculated as described in Material and Methods to assess animal cleanliness and production hygiene.

| Farm | Prevalence *Lm*<br>animals | Score animals | Prevalence *Lm*<br>food trough | Score food<br>trough | Prevalence *Lm*<br>water trough | Score water<br>trough | Prevalence *Lm*<br>bedding | Score bedding | Prevalence *Lm*<br>milk filter | Score milking<br>tank | Prevalence *Lm*<br>milk parlor | Score milking<br>parlor |
| --- | --- | --- | --- | --- | --- | --- | --- | --- | --- | --- | --- | --- |
| CATTLE A | 0.00 | 2.1 | 0.0 | 3 | 0 | 3 | 11.1 | 1 | 0 | 3 | 0 | 2 |
| CATTLE B | 9.38 | 1.9 | 0.0 | 3 | 0 | 3 | 22.2 | 3 | 0 | 2 | 0 | 2 |
| CATTLE C | 3.13 | 2.1 | 11.1 | 3 | 0 | 3 | 0.0 | 2 | 0 | 3 | 0 | 3 |
| CATTLE D | 0.00 | 2.5 | 0.0 | 2 | 0 | 2 | 0.0 | 1 | 0 | 1 | 0 | 1 |
| CATTLE E | 4.17 | 4.1 | 0.0 | 3 | 0 | 2 | 0.0 | 1 | 0 | 3 | 0 | 1 |
| CATTLE F | 0.00 | 1.7 | 0.0 | 3 | 0 | 2 | 0.0 | 1 | 0 | 3 | 0 | 3 |
| CATTLE G | 14.58 | 3.0 | 66.7 | 3 | 0 | 1 | 0.0 | 2 | 0 | 2 | 0 | 3 |
| CATTLE H | 1.04 | 1.8 | 0.0 | 3 | 11.1 | 2 | 0.0 | 3 | 0 | 3 | 0 | 2 |
| CATTLE I | 10.42 | 1.5 | 33.3 | 3 | 0 | 2 | 22.2 | 3 | 0 | 2 | 0 | 3 |
| CATTLE J | 0.00 | 2.2 | 0.0 | 3 | 0 | 1 | 0.0 | 3 | 0 | 1 | 0 | 2 |
| (P> 0.05) Spearman's rho | 0.056 |  | 0.214 |  | 0.063 |  | 0.337 |  | n.c. |  | n.c. |  |
| SHEEP A | 0,0 | 3 | 0,0 | 3 | 0,0 | 2 | 0 | 3 | 0,0 | 3 | 0,0 | 3 |
| SHEEP B | 28.1 | 3 | 0,0 | 2 | 0,0 | 1 | 0 | 1 | 44.4 | 3 | 0,0 | 3 |
| SHEEP C | 1,0 | 2.5 | 0,0 | 2 | 0,0 | 2 | 11.1 | 2 | 0,0 | 2,4 | 0,0 | 2,4 |
| SHEEP D | 0,0 | 2 | 0,0 | 3 | 0,0 | 3 | 0 | 3 | 0,0 | 3 | 0,0 | 3 |
| SHEEP E | 0,0 | 2 | 0,0 | 2 | 0,0 | 2 | 0 | 2 | 0,0 | 2 | 0,0 | 2 |
| (P> 0.05) Spearman's rho | 0.53 |  | n.c. |  | n.c. |  | 0.186 |  | 0.395 |  | n.c. |  |
| GOAT A | 1.0 | 1 | 0,0 | 2 | 0,0 | 2 | 0,0 | 2 | 0,0 | 3 | 0,0 | 3 |
| GOAT B | 0,0 | 1 | 0,0 | 2 | 0,0 | 1 | 0,0 | 3 | 0,0 | 3 | 0,0 | 3 |
| GOAT C | 0,0 | 1 | 0,0 | 2 | 0,0 | 3 | 0,0 | 3 | 0,0 | 3 | 0,0 | 3 |
| GOAT D | 0,0 | 1 | 0,0 | 3 | 0,0 | 3 | 0,0 | 3 | 0,0 | 3 | 0,0 | 3 |

n.c., not calculated (prevalence = 0)

